# A multi-trait meta-analysis with imputed sequence variants reveals twelve QTL for mammary gland morphology in Fleckvieh cattle

**DOI:** 10.1101/030981

**Authors:** Hubert Pausch, Reiner Emmerling, Hermann Schwarzenbacher, Ruedi Fries

## Abstract

**Background:** The availability of whole-genome sequence data from key ancestors provides an exhaustive catalogue of polymorphic sites segregating within and across cattle breeds. Sequence variants from key ancestors can be imputed in animals that have been genotyped using medium- and high-density genotyping arrays. Association analysis with imputed sequences, particularly if applied to multiple traits simultaneously, is a very powerful approach to revealing candidate causal variants underlying complex phenotypes.

**Results:** We used whole-genome sequence data from 157 key ancestors of the German Fleckvieh population to impute 20 561 798 sequence variants in 10 363 animals that had (partly imputed) array-derived genotypes at 634 109 SNP. The imputed sequence data were enriched for rare variants. Association studies with imputed sequence variants were performed using seven correlated udder conformation traits as response variables. The calculation of an approximate multi-trait test statistic enabled us to detect twelve major QTL (P<2.97 × 10^−9^) controlling different aspects of mammary gland morphology. Imputed sequence variants were the most significantly associated at eleven QTL, whereas the top association signal at a QTL on BTA14 resulted from an array-derived variant. Seven QTL were associated with multiple phenotypes. Most QTL were located in non-coding regions of the genome in close neighborhood, however, to plausible candidate genes for mammary gland morphology (*SP5, GC, NPFFR2, CRIM1, RXFP2, TBX5, RBM19, ADAM12*).

**Conclusions:** Association analysis with imputed sequence variants allows QTL characterization at maximum resolution. Multi-trait approaches can reveal QTL that are not detected in single-trait association studies. Most QTL for udder conformation traits were located in non-coding elements of the genome suggesting regulatory mutations to be the major determinants of variation in mammary gland morphology in cattle.

## Background

Genome-wide association studies (GWAS) using dense SNP facilitated the identification of quantitative trait loci (QTL) for numerous phenotypes. However, the extensive linkage disequilibrium in cattle populations typically resulted in large QTL intervals and made the identification of underlying genes and variants often impossible. Moreover, the current genotyping arrays interrogate only a limited number of polymorphic sites that are primarily located in non-coding regions of the genome [1].

The availability of whole-genome sequences makes it possible to compile an exhaustive catalogue of polymorphic sites segregating within and across cattle populations [2],[3]. Obtaining genome-wide sequence data for a large number of animals is still too costly. However, a relatively low number of sequenced key ancestors may serve as a reference to impute sequence variants in any animal with dense genotyping data [4],[5]. Association studies with imputed sequence variants can then pinpoint candidate causal variants controlling complex trait variation [6],[3].

Computationally efficient algorithms facilitate to perform association studies in thousands of individuals that have been genotyped at millions of polymorphic sites (*e.g,* [7],[8]). Such association studies in cattle are typically performed within breed on a trait-by-trait basis and by testing one variant at a time. Association analyses involving multiple phenotypes may be more powerful in identifying QTL for complex traits, particularly with causal variants affecting multiple correlated phenotypes [9]. However, multivariate association testing of millions of sequence variants with a large number of phenotypes is computationally challenging [9]. An approximate multi-trait test statistic allows to efficiently combine the results of multiple separately performed association studies and thereby increases the power to identify trait-associated variants [10].

Udder conformation traits are routinely recorded in cattle populations during the appraisal of first-crop daughters of test bulls. Phenotypes for such traits are an important source of information because mammary gland morphology is highly correlated with mastitis susceptibility and productive life span [11],[12],[13]. Although the definitions of udder conformation traits may vary across breeds, phenotypic records typically describe the teat morphology and placement and the overall shape of the mammary gland. The heritability of such traits is relatively high ranging from 0.14 to 0.42 and most traits describing mammary gland morphology are correlated with each other [11],[14].

Here we present results that are based on the imputation of whole-genome sequence variants in more than 10 000 Fleckvieh animals that have been genotyped using dense SNP arrays. We performed association studies with more than 16 million sequence variants using seven highly correlated udder conformation traits as response variables. The multi-trait meta-analysis enabled us to detect twelve major QTL controlling different aspects of mammary gland morphology in Fleckvieh cattle.

## Methods

### Animal ethics statement

DNA of artificial insemination bulls was prepared from semen samples that were collected by approved commercial artificial insemination stations as part of their regular breeding and reproductive measures in cattle industry. No ethical approval was required for this study.

### Genotypes of the target population

All animals were genotyped using medium- and high-density SNP arrays. The high-density dataset consisted of 3545 Fleckvieh animals that had been genotyped with the Illumina BovineHD Bead chip comprising 777 962 SNP. The medium-density dataset consisted of 7073 Fleckvieh animals that had been genotyped with the Illumina BovineSNP50 Bead chip (version 1 and version 2) comprising approximately 54 000 SNP. The chromosomal position of the SNP corresponded to the UMD3.1 assembly of the bovine genome [15]. Mitochondrial, X-chromosomal, Y-chromosomal and SNP with unknown chromosomal position were not considered for further analyses. Quality control (call-rate per SNP and per individual higher than 90%, no deviation from the Hardy-Weinberg equilibrium (P>0.0001), minor allele frequency (MAF) above 0.5%, no pedigree conflicts) was carried out separately for each dataset as detailed by Pausch et al. [16]. After quality control, the medium-density genotypes were imputed to higher density using a combination of *Beagle* [17] and *Minimac* [18] as detailed by Pausch et al. [5]. Only SNP with a MAF above 0.5% were retained after imputation. The final array-derived dataset consisted of 10 363 animals with (partly imputed) genotypes at 634 109 autosomal SNP.

### Generation of sequence data

We used whole-genome sequence data from 263 animals representing ten cattle breeds, among them 157 Fleckvieh animals. Most of the sequenced animals were key-ancestors for their breeds [19]. The generation and analysis of whole-genome sequence data is detailed by Pausch et al. [20]. Single nucleotide along with short insertion and deletion polymorphisms were genotyped in all 263 sequenced animals simultaneously using the multi-sample approach implemented in the *mpileup* module of *SAMtools* [21] and in a variant calling pipeline described by Jansen et al. [2]. A total of 25 426 490 polymorphic sites were identified. The functional effects of the sequence variants were analyzed using the annotation of the UMD3.1 assembly of the bovine genome [22] and as detailed by Jansen et al. [2].

### Imputation of sequence variants

The imputation reference panel consisted of 157 sequenced Fleckvieh animals with 20 561 798 autosomal variants. Haplotypes were inferred using *Beagle* [17] and served as reference to impute genotypes for 20 561 798 variants in 10 363 target animals with (partly imputed) genotypes at 634 109 SNP (see above) using *Minimac* [18].

### Phenotypes for association testing

Response variables for association testing were daughter yield deviations (DYD) for seven udder conformation traits (teat thickness, teat length, teat position, udder depth, central ligament, fore udder attachment and fore udder length). Only DYD with an accuracy greater than 0.7 were considered for association testing. Depending on the trait, the number of animals with phenotypes ranged from 5470 to 7159 (Table 1). In addition, association testing of 682 047 imputed sequence variants located on BTA14 was carried out in 6838 animals using DYD for height at the sacral bone as response variables.

**Table 1:** Characteristics of seven udder conformation traits

| Phenotype (and abbreviation) | Number of animals with phenotypes | Distribution of the DYD |  |  | Accuracy of the DYD |  |  |
| --- | --- | --- | --- | --- | --- | --- | --- |
| | | min | max | mean (± sd) | min | max | mean (± sd) |
| Udder depth (UD) | 7110 | -2.22 | 2.75 | 0.11 (±0.64) | 0.71 | 0.99 | 0.91 (±0.04) |
| Teat thickness (TT) | 7108 | -2.32 | 2.17 | 0.04 (±0.52) | 0.71 | 0.99 | 0.91 (±0.04) |
| Teat length (TL) | 7159 | -2.61 | 2.52 | -0.02 (±0.66) | 0.71 | 0.99 | 0.93 (±0.04) |
| Teat placement (TP) | 7108 | -1.92 | 2.50 | 0.16 (±0.55) | 0.71 | 0.99 | 0.91 (±0.04) |
| Fore udder length (FUL) | 7034 | -2.03 | 2.15 | 0.21 (±0.51) | 0.71 | 0.99 | 0.88 (±0.04) |
| Central ligament (CL) | 6923 | -2.24 | 2.32 | 0.11 (±0.57) | 0.71 | 0.99 | 0.85 (±0.05) |
| Fore udder attachment (FUA) | 5470 | -2.73 | 2.48 | 0.13 (±0.65) | 0.71 | 0.99 | 0.87 (±0.05) |

### Single-trait genome-wide association studies

We considered 16 816 809 imputed sequence variants with a MAF above 0.5% for the GWAS. The imputed sequence variants were tested for association with each trait in turn using a two-step variance components-based approach as implemented in the *EMMAX* software tool [7]: in a first step, the polygenic and error variances were estimated using following variance component model: 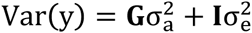, where **G** is the realized genomic relationship matrix of the 10 363 animals built using genotypes of 634 109 autosomal SNP following VanRaden’s approach [23], **I** is an identity matrix, 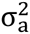 is the polygenic variance and 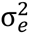 is the error variance. In a second step, the allele substitution effect (b) is obtained from a generalized linear regression model: **y** = **μ** + **x**b + **e**, where μ is the intercept, **x** is a vector of expected allele dosages and **e** is a vector of random residual deviates with variance 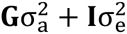. Inflation factors were calculated using the *estlamdba()-function* of *GenABEL* [24]. Sequence variants with P<2.97 × 10^−9^ were considered as significantly associated (5% Bonferroni-corrected significance threshold for 16 816 809 independent tests). An analysis conditional on the most significantly associated variant was carried out by taking the expected allele dosages of the top variant as covariates in the linear regression model (see above).

### Multi-trait meta-analysis

An approximate multi-trait test statistic was calculated for 16 816 809 imputed sequence variants using 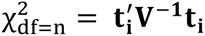, where n is the number of traits, **t_i_** is a n×1 vector of t-values 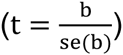 of the i^th^ SNP and **V^−1^** is the inverse of the n×n correlation matrix of t-values [10]. The correlation matrix **V** was constructed from t-values of 16 816 809 imputed sequence variants. Sequence variants with P_META_<2.97 × 10^−9^ were considered as significantly associated (see above).

## Results

More than 20 million sequence variants were imputed in 10 363 animals that had (partly imputed) array-derived genotypes for 634 109 SNP (Figure 1A). The allele frequency distribution of the imputed sequence (SEQ) variants differed from the distribution of the array-derived variants. Variants from medium-(50K) and high-density (700K) SNP arrays were almost uniformly distributed across different MAF classes whereas the imputed sequence variants were enriched for low-frequency classes (Figure 1B). The proportions of variants with MAF below 0.05 were 10.66%, 9.04% and 40.52% and the average MAF was 0.249, 0.260 and 0.145 for the 50K, 700K and SEQ dataset, respectively.

**Figure 1.**
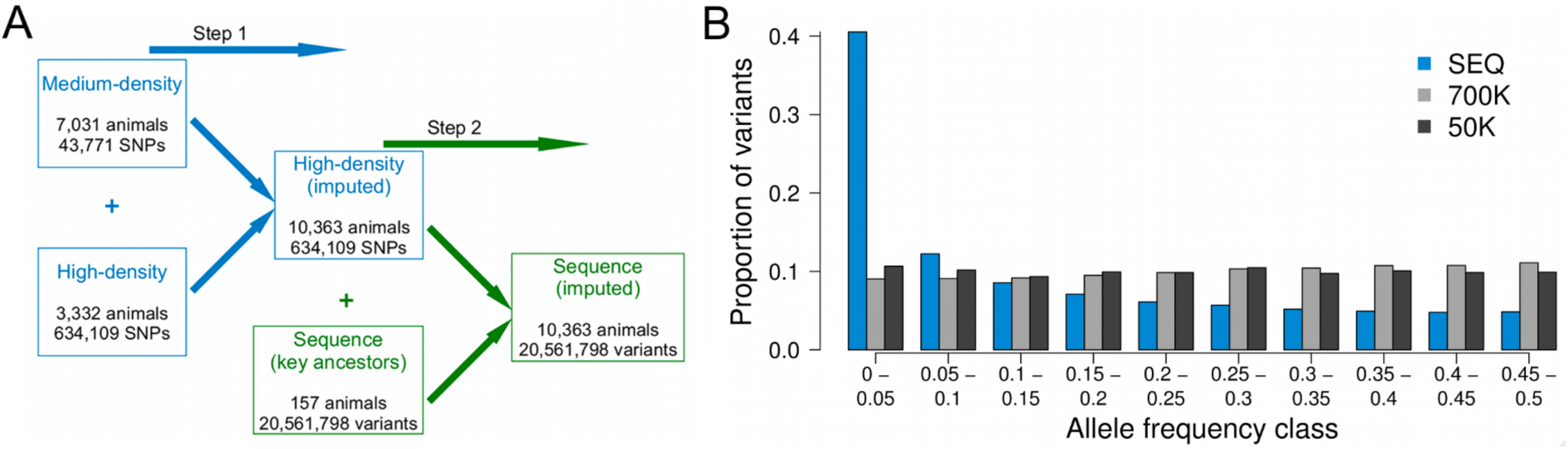
Imputation of sequence variants: Overview of the stepwise imputation of 20 561 798 sequence variants in 10 363 Fleckvieh animals (A). Allele frequency distribution of imputed and array-derived variants (B). Blue and shades of grey represent the proportion of imputed sequence (SEQ) and array-derived (50K, 700K) variants, respectively, for ten allele frequency classes.

### Association studies with udder conformation traits

Association studies of 16 816 809 imputed sequence variants with MAF above 0.5% were performed with DYD for seven udder conformation traits (Table 1). The inflation factors of the association studies ranged from 1.038 (central ligament) to 1.081 (udder depth) with an average inflation factor of 1.061 indicating proper control of population stratification. From zero (fore udder attachment) to five (fore udder length and teat placement) QTL were detected per trait [See Additional File 1]. Correlation coefficients among the seven traits were calculated with the signed t-values (*i.e.,* allele substitution effect divided by its standard error, Figure 2A). The highest correlations were observed between udder depth and fore udder attachment (r=0.47), teat length and teat thickness (r=0.46) and central ligament and fore udder attachment (r=0.37).

**Figure 2.**
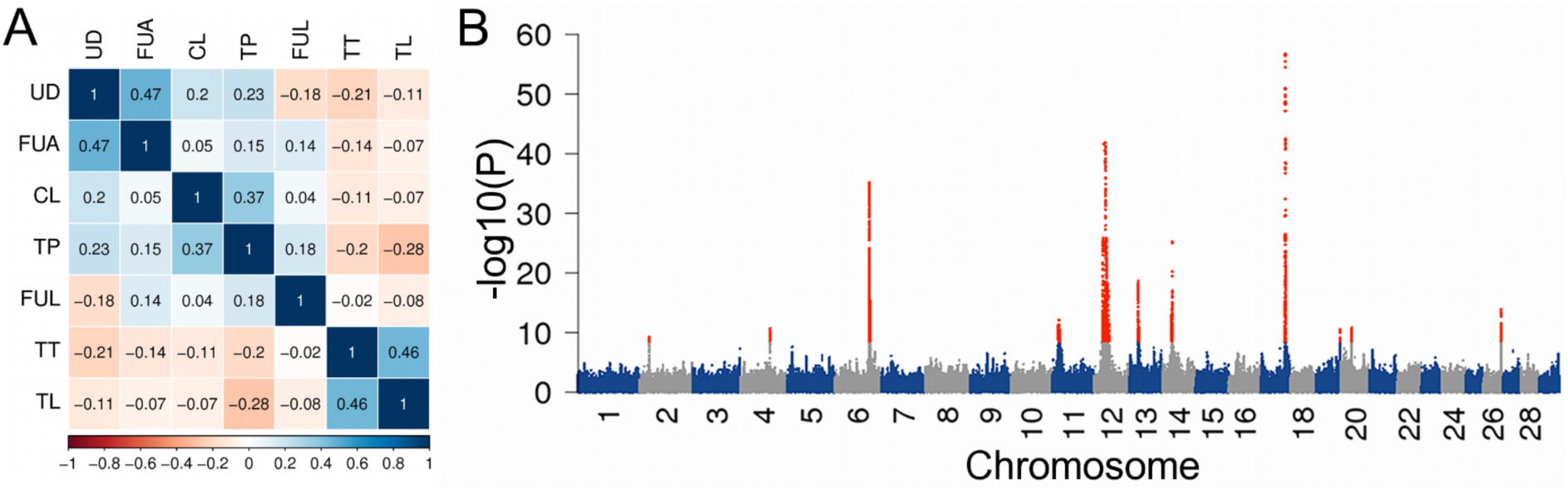
Identification of twelve QTL for mammary gland morphology: Correlations between seven udder conformation traits (A). The abbreviations of the traits are listed in Table 1. Shades of blue and red indicate positive and negative correlation coefficients, respectively. Manhattan plot representing the association of 16 816 809 imputed sequence variants in the multi-trait meta-analysis (B). Red color represents variants with P_META_<2.9 × 10^−9^.

A meta-analysis of the seven single-trait association studies revealed eleven QTL (P_META_<2.9 × 10^−9^) located on eleven chromosomes (Figure 2B, Table 2). Imputed sequence variants were the most significantly associated variants for ten QTL, whereas the top association signal for the QTL on BTA14 resulted from an array-derived SNP. The multi-trait meta-analysis revealed a QTL on BTA2 (P_META_=6.1 × 10^−10^) that was not detected in the single-trait association studies. Closer inspection revealed association of the QTL on BTA2 with fore udder attachment (P_SINGLE_=1.13 × 10^−6^), teat thickness (P_SINGLE_=2.68 × 10^−4^), fore udder length (P_SINGLE_=4.37 × 10^−3^) and udder depth (P_SINGLE_=5.01 × 10^−3^), although above the Bonferroni-corrected threshold of genome-wide significance. Four QTL on BTA14, BTA21, BTA27 and BTA29 that were associated with teat placement, udder depth, teat length and udder depth, respectively, were not identified in the multi-trait meta-analysis [See Additional File 1]. Closer inspection of the four regions revealed association in the multi-trait meta-analysis (7.91 × 10^−8^<P_META_<1.98 × 10^−6^), however not at genome-wide significance. Seven QTL were associated with multiple aspects of mammary gland morphology (Table 2, Figure 3). The QTL on BTA17 was associated with four phenotypes. Two and four QTL were associated with three and two udder conformation traits, respectively.

**Table 2:** Position of twelve QTL for udder morphology

| Chr | Position | MAF | $P_{\text{META}}$ | Candidate gene(s) |
| --- | --- | --- | --- | --- |
| 2 | 25 887 326 | 0.25 | $6.1 \times 10^{-10}$ | *SP5* |
| 4 | 75 817 266 | 0.24 | $1.9 \times 10^{-11}$ | - |
| 6 | 88 723 742 | 0.47 | $7.4 \times 10^{-36}$ | *GC, NPFFR2* |
| 6* | 90 366 765 | 0.21 | $8.5 \times 10^{-16}$ | *RASSF6* |
| 11 | 18 757 907 | 0.28 | $7.5 \times 10^{-13}$ | *CRIM1* |
| 12 | 29 248 113 | 0.03 | $1.6 \times 10^{-42}$ | *RXFP2* |
| 13 | 22 699 039 | 0.29 | $2.2 \times 10^{-19}$ | - |
| 14 | 25 015 640 | 0.12 | $6.2 \times 10^{-26}$ | *PLAG1* |
| 17 | 62 694 032 | 0.25 | $2.2 \times 10^{-57}$ | *TBX5, RBM19* |
| 19 | 61 493 292 | 0.35 | $3.4 \times 10^{-11}$ | - |
| 20 | 27 355 591 | 0.15 | $1.8 \times 10^{-11}$ | - |
| 26 | 46 058 305 | 0.34 | $1.3 \times 10^{-14}$ | *ADAM12* |

The position of the QTL corresponds to the UMD3.1 assembly of the bovine genome. The asterisk marks a QTL that was detected in the conditional analysis. MAF: minor allele frequency.

**Figure 3.**
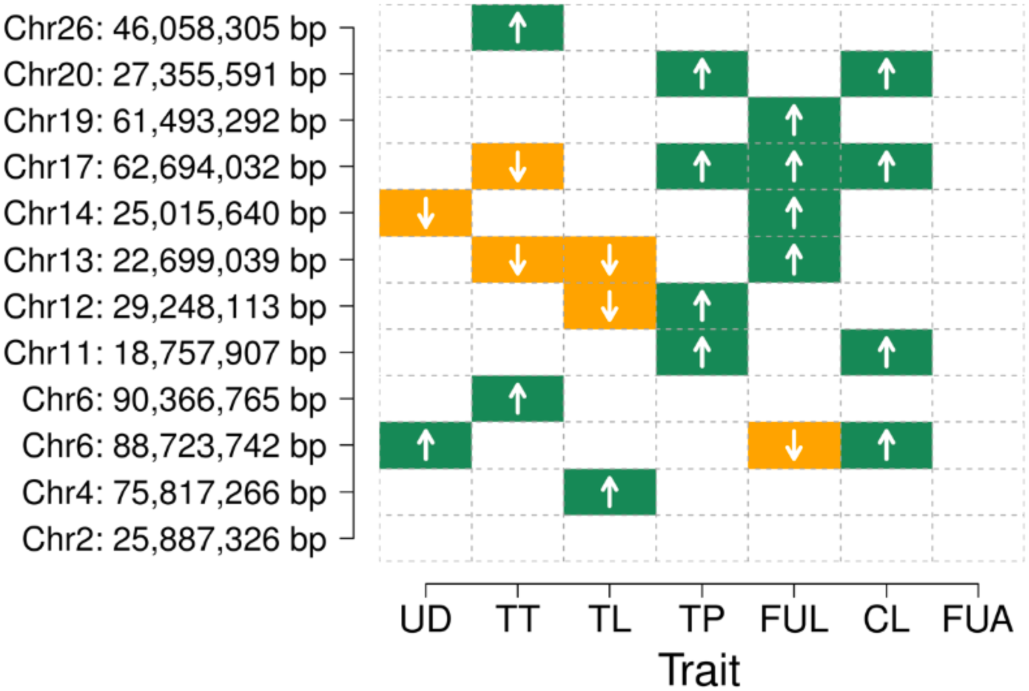
The effect of twelve QTL on seven udder conformation traits: Green and orange colour represents positive and negative phenotype × genotype associations, respectively, of twelve QTL in seven single-trait association studies. The abbreviations of the traits are listed in Table 1. Only associations with P<2.9 × 10^−9^ (Bonferroni-corrected significance threshold for 16 816 809 tests) are shown.

To test if a QTL was completely tagged by the top variant, the most significantly associated variant was fitted as a covariate in the single-trait GWAS model and the multi-trait test statistic was re-calculated. The conditional analysis revealed that the associated region on BTA6 consists of two distinct closely neighboring QTL [See Additional File 2]. The QTL located at 88 723 742 bp was associated with udder depth, central ligament and fore udder length whereas the second QTL located at 90 366 765 bp was associated with teat thickness (Table 2, See Additional File 2). There were no significant associations detected for all other QTL when the association analysis was conditioned on the respective top SNP.

### Gene content of twelve QTL-regions

To detect positional and functional candidate genes controlling mammary gland morphology, we examined the gene content of twelve QTL regions identified by the meta-analysis [See Additional File 3]. However, there were no annotated genes within a 300 kb interval on either side of the top association signal on BTA4, BTA13, BTA19 and BTA20, precluding the identification of positional candidate genes.

Seventy-seven variants with P_META_<1 × 10^−8^ were located within a 100 kb intergenic segment on BTA2 [See Additional File 3]. The top association signal (P_META_=6.11 × 10^−10^) resulted from a variant (25 887 326 bp) that resided 86 kb downstream of the stop codon of *MYO3B* encoding myosin IIIB and 111 kb upstream of the translation start of *SP5* encoding the Sp5 transcription factor.

A QTL-region on BTA6 encompassed 1199 variants with P_META_<1 × 10^−14^ located within a 450 kb segment (88 613 408bp – 89 062 806 bp) [See Additional File 3]. Two annotated genes reside within this interval: the group specific component-encoding gene *GC* and the neuropeptide FF receptor 2-encoding gene *NPFFR2.* The top variant (88 723 742 bp; P_META_=7.44×10^−36^) was located in the first intron of *GC.* However, the most significantly associated coding variant (*NPFFR2*:pE389K; P_META_=186 × 10^−16^) was considerably less significantly associated than the non-coding top variant. A second QTL on BTA6 was located 6 kb downstream of the stop codon of *RASSF6* encoding the Ras association (RalGDS/AF-6) domain family member 6 [See Additional File 3].

Eighty-one variants located within a 1.5 Mb interval on BTA11 (18 546 414 bp – 20 048 201 bp) had P_META_-values <3.1 × 10^−10^ [See Additional File 3]. This segment encompasses twelve annotated transcripts/genes. The top variant (18 757 907 bp; P_META_=7.47×10^−13^) was located 77 kb upstream of the translation start of *CRIM1,* encoding cysteine rich transmembrane BMP regulator 1. Two highly significantly associated coding variants (P_META_<1.58 × 10^−10^) in *CRIM1* (p.R540K) and *PRKD3* (p.R864H) were in high linkage disequilibrium (r^2^>0.88) with the top non-coding variant.

One hundred and twelve variants with P_META_<1 × 10^−30^ were located in a 4.59 Mb interval (25 693 051 bp – 30 288 956 bp) on BTA12 [See Additional File 3]. The top association signal (P_META_=1.58×10^−42^) resulted from an intronic variant (29 248 113 bp) in *RXFP2,* encoding relaxin/insulin-like family peptide receptor 2. None of the highly significantly associated variants were located in the coding region of an annotated gene.

The QTL on BTA14 was in a genomic region that controls growth-related traits in cattle [25],[26] and other species [27]. The top variant (BovineHD1400007259, 25 015 640 bp, P_META_=6.23 × 10^−26^) was located 6 kb upstream of the translation start of *PLAG1* (pleiomorphic adenoma gene 1) [See Additional File 3]. We performed an association analysis with DYD for height at the sacral bone to test if the mammary gland morphology QTL is also associated with stature. The association analysis revealed that BovineHD1400007259 was also the most significantly associated variant for height at the sacral bone (P=1.07 × 10^−52^) [See Additional File 4]. The allele that increases body height was associated with an increased udder base. Eighty sequence variants with P_META_<1.87 × 10^−25^ were located within a 171 kb intergenic segment (62 667 848 bp – 62 838 591 bp) on BTA17 [See Additional File 3]. The top variant (62 694 032 bp; P_META_=2.23 × 10^−57^) was located 193 kb upstream of the translation start of *RBM19,* encoding RNA binding motif protein 19, and 113 kb downstream of the translation end of *TBX5,* encoding T-box 5 transcription factor.

A QTL on BTA26 was associated with teat thickness. Thirty-nine variants with P_META_-values <2.01 × 10^−13^ were located in the third intron of *ADAM12,* encoding ADAM metallopeptidase domain 12 [See Additional File 3]. The top variant (P_META_=1.29 × 10^−14^) was located at 46 058 305 bp. There were no coding variants significantly associated with mammary gland morphology.

## Discussion

Association studies with 16 816 809 imputed sequence variants were carried out in up to 7159 Fleckvieh animals using DYD for seven udder conformation traits as response variables. The calculation of a multi-trait test statistic enabled us to detect twelve major QTL for mammary gland morphology.

Sequence variants were extrapolated in 10 363 animals using a two-step genotype imputation approach. Initially, the animals with medium-density genotypes were imputed to higher density using 3332 reference animals that had been genotyped with a high-density genotyping array. In a second step, the (partly imputed) high-density genotypes were imputed to sequence level using sequence variants of 157 key ancestors as a reference. Low-frequency variants occurred more often among the imputed than the array-derived variants. This agrees with previous findings in cattle [6] and other species [28]. Since the imputation of rare variants is error prone [5],[3],[29], we retained only variants with a MAF above 0.5% for the association studies. Moreover, to take imputation uncertainty into account [30], we used the expected allele dosages instead of the most likely genotypes as explanatory variables in the GWAS. Thus, our association analyses should not be flawed due to inaccurately imputed sequence variants.

To eliminate false positive association signals due to population stratification, the genomic relationship based on genome-wide marker data was considered when carrying out seven separate association studies. Low inflation factors (1.038 - 1.081) evidence the success of this corrective measure. Combining the results of the seven separate association studies by calculating an approximate multi-trait test statistic enabled us to reveal twelve major QTL for mammary gland morphology. Among them was a QTL on BTA2 that was not detected in the single-trait studies, showing the enhanced capacity of multi-trait approaches for detecting QTL, particularly when the phenotypes are correlated [9],[10],[31]. The QTL on BTA2 was associated with four correlated traits, although not on a genome-wide scale. We corroborate the findings of O’Reilly et al. [31] by showing again that QTL which cannot be detected at a genome-wide significance level in single-trait GWAS can be uncovered in a multi-trait approach. The joint association analysis of multiple phenotypes might be a more powerful approach to detect QTL underlying correlated traits than the multi-trait test statistic applied in our study [9]. However, phenotypic records were incomplete for some animals of the present study precluding multivariate association testing without relying on algorithms for estimating missing phenotypes. Four QTL were detected in single-trait association studies but were not formally identified in the multi-trait metaanalysis. However, the corresponding P_META_-values were only slightly above the Bonferroni-corrected genome-wide significance threshold. Adjusting for multiple testing using Bonferroni-correction assumes the individual tests to be independent from each other. Due to the low effective population size and high linkage disequilibrium, this assumption does not hold for genome-wide association studies in livestock populations. The Bonferroni-correction method is prone to an over-correction, particularly in association studies involving millions of sequence variants [32],[33], resulting in a reduced power of such studies.

Association studies with udder conformation traits had been carried out in several cattle breeds using dense marker maps. Flury et al. [34] identified two QTL for udder length and teat diameter in the Brown Swiss cattle breed on BTA6 located at 88.92 Mb and 90.37 Mb, respectively. We also identified two QTL on BTA6 at 88.72 Mb and 90.37 Mb, corroborating the crucial role of both regions for mammary gland morphology in cattle. In our study, the QTL on BTA6 were associated with teat thickness and fore udder length, central ligament and udder depth. Hiendleder et al. identified a QTL for udder conformation traits in the Holstein breed on BTA6 at 88 cM [35], which most likely corresponds to the highly significantly associated region(s) identified in our study. Another QTL affecting mammary gland morphology in cattle has been reported on BTA17, close to *TBX3, TBX5* and *RBM19* [36],[34]. The corresponding human chromosome segment is involved in ulnar mammary syndrome [37]. We also detected a highly significantly associated QTL at that position. The top variant was only 3 kb distant from the top association signal reported in the Brown Swiss breed [34], indicating that a common variant might control udder traits in both breeds. The position of the QTL suggests the involvement of regulatory variants in shaping the mammary gland. An improved functional annotation of the bovine genome [38] and multi-breed association studies with imputed sequence variants may reveal causal mutations for such QTL.

Imputed sequence variants were the most significantly associated at eleven QTL, demonstrating that a better mapping resolution can be obtained with whole-genome sequence data than with array-derived genotypes. However, the top association signal at a QTL on BTA14 resulted from an array-derived variant (BovineHD1400007259). Interestingly, BovineHD1400007259 is among eight candidate causative trait variants for a QTL that affects bovine stature by modulating the expression of *PLAG1* [25]. In our study, BovineHD1400007259 was also the top variant for body height indicating pleiotropic effects on stature and udder traits. The allele that increases height was also associated with an increased udder base. Udder depth, *i.e.* the interspace between the ankle and the udder base, is visually examined during the appraisal of first crop daughters of artificial insemination bulls. It is possible that the interspace is overestimated in tall animals. Thus the association of the *PLAG1* region with udder depth may reflect phenotypic variation in body size rather than true effects on mammary gland morphology. The absence of the association when udder depth was conditioned on body height further supports this assumption [See Additional File 5].

Our association study revealed new candidate genes for mammary gland morphology in cattle. A QTL on BTA2 was located close to *SP5,* encoding the transcription factor Sp5. Since Sp5 is a downstream target of the Wnt signaling [39], our findings provide additional evidence for the crucial role of the Wnt signaling pathway for mammary gland development in cattle [36]. A QTL on BTA6 was located in the region of *GC* and *NPFFR2,* two genes that had been implicated in mastitis susceptibility in cattle [40],[41],[42]. In our study, the QTL was associated with udder depth and central ligament. Udder depth, central ligament and mastitis susceptibility are negatively correlated traits [13] suggesting that the QTL on BTA6 might contribute to the unfavorable genetic correlation between a deep udder base and udder health. A QTL on BTA11 is located close to *CRIM1,* encoding a transmembrane protein, which contains an insulin-like growth factor-binding domain [43]. The QTL-region on BTA12 contains *RXFP2* that encodes relaxin/insulin-like family peptide receptor 2. The QTL on BTA26 is located in an intronic region of *ADAM12,* encoding ADAM metallopeptidase domain 12 that interacts with insulin-like growth factor-binding proteins [44]. Such findings suggest a crucial role of insulin-like growth factors and insulin-like growth factor-binding proteins during mammary gland development [45],[46].

Most of the QTL identified in our study reside in non-protein coding regions of the genome indicating a crucial role of regulatory mutations for phenotypic variation in mammary gland morphology in cattle. Pinpointing causal mutations in non-coding elements is notoriously difficult since the annotation of the bovine genome is often flawed due to assembly problems or gaps in the reference sequence [22]. Moreover, regulatory elements in the bovine genome are poorly characterized. Thus we did not attempt to identify candidate causal variants for QTL in the present study. However, an improved functional annotation of the bovine genome is expected to facilitate a more precise characterization of regulatory QTL in the future [38].

## Conclusions

Association analysis with imputed sequence variants allows for QTL characterization at maximum resolution. Variants that affect multiple correlated traits are most efficiently uncovered by their simultaneous analysis using a multi-trait test statistic. Our study revealed twelve QTL controlling different aspects of mammary gland morphology in the German Fleckvieh population. The positions of the QTL suggest variants of regulatory elements to be major determinants for phenotypic variation in mammary gland morphology in cattle.

## List of abbreviations

DYD: daughter yield deviation; MAF minor allele frequency; GRM: genomic relationship matrix; QTL: quantitative trait locus; SNP: single nucleotide polymorphism

## Authors’ contribution

HP and RF conceived, designed and performed the experiments. RE and HS contributed pedigree, phenotype and genotype data. HP and RF wrote the manuscript. All authors have read and approved the final manuscript.

## Acknowledgements

The generation of sequence data of Fleckvieh animals was funded by the German Federal Ministry of Education and Research (BMBF) within the AgroClustEr “Synbreed – Synergistic plant and animal breeding” (grant ids 0315527B, 0315528A). We acknowledge the Arbeitsgemeinschaft Süddeutscher Rinderzüchter e.V., the Arbeitsgemeinschaft österreichischer Fleckviehzüchter and ZuchtData EDV Dienstleistungen GmbH for providing genotype data. We thank Qualitas AG (CH-Zug) and the Swiss Cattle Breeder Association for funding the sequencing of original Simmental animals.

---

**Additional file 1.**
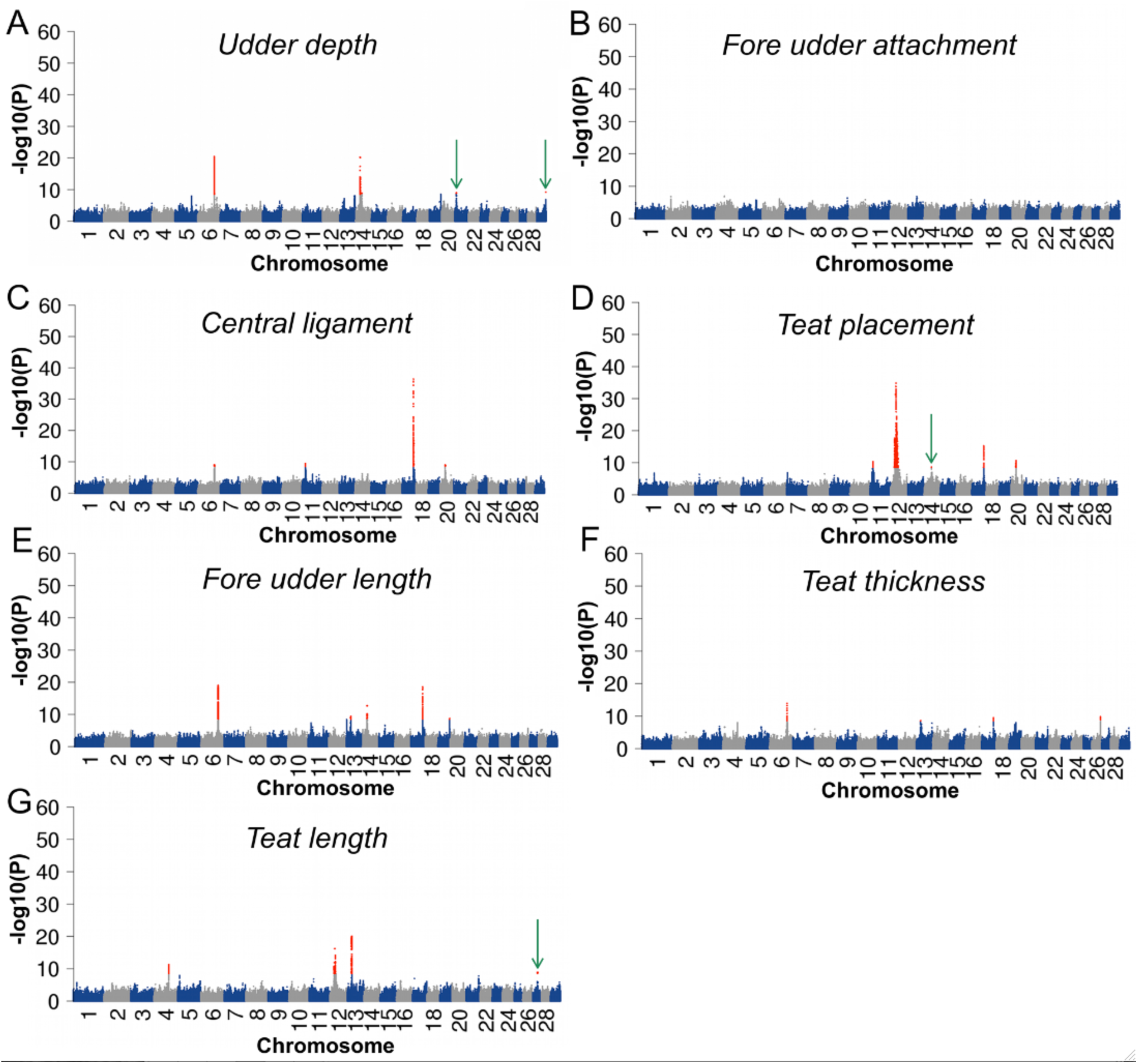
Manhattan plots representing the association of 16 816 809 imputed sequence variants with seven udder conformation traits. Red dots represent variants with P_SINGLE_<2.9 × 10^−9^. Green arrows highlight four QTL that were not detected in the multitrait meta-analysis.

**Additional file 2.**
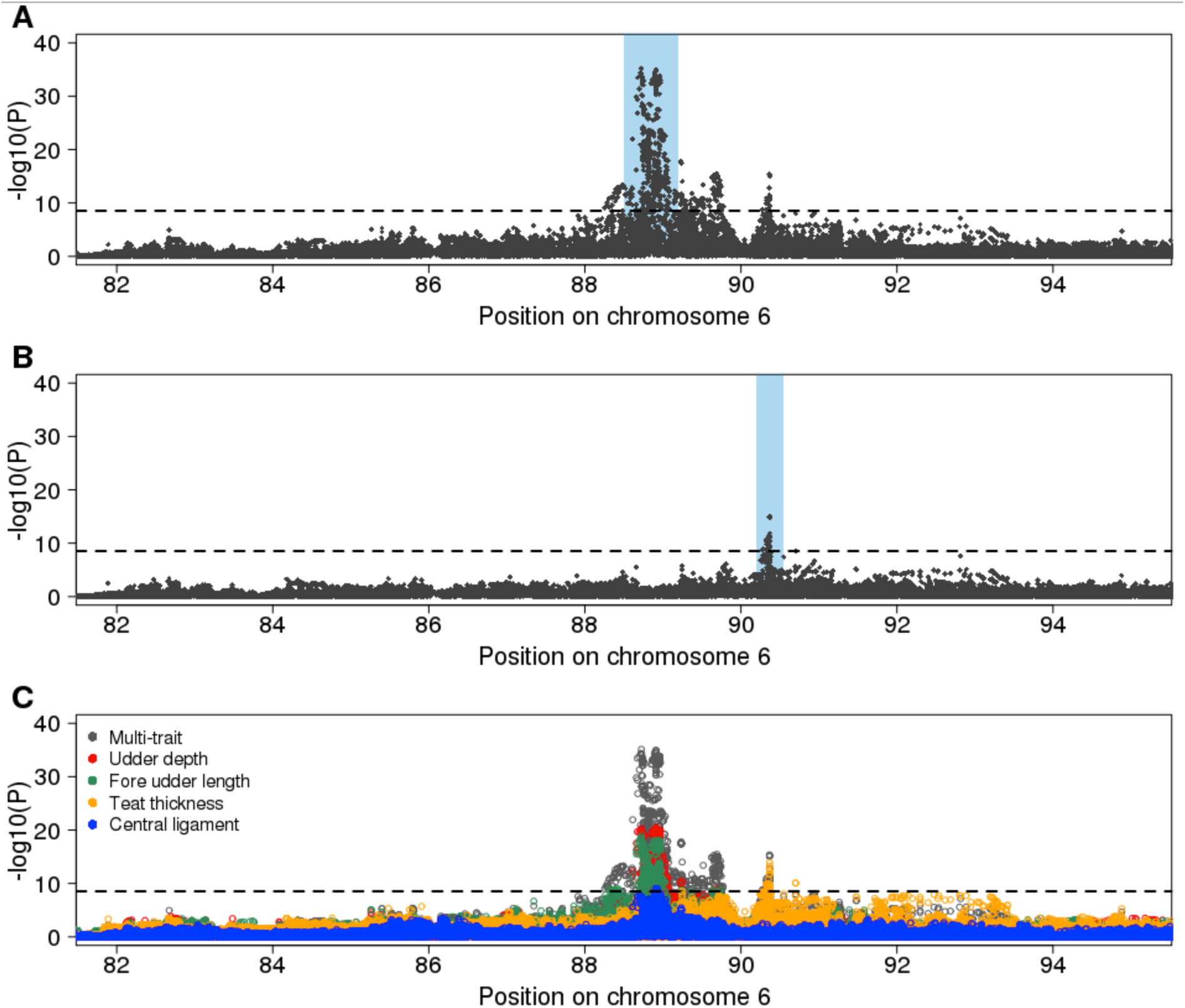
Detailed view of the QTL on BTA6. Manhattan plots representing the association of 91 355 imputed sequence variants located in an interval from 82 000 000 bp – 95 000 000 bp on chromosome 6 with different aspects of mammary gland morphology. The dotted line represents the Bonferroni-corrected significance threshold. The multi-trait meta-analysis identified a QTL located at 88 723 742 bp (blue rectangle) (A). Another QTL located at 90 366 765 bp (blue rectangle) was identified after conditioning on the top variant (B). The QTL at 88 723 742 bp is associated with udder depth, fore udder length and central ligament whereas the QTL at 90 366 765 bp is associated with teat thickness (C).

**Additional file 3.**
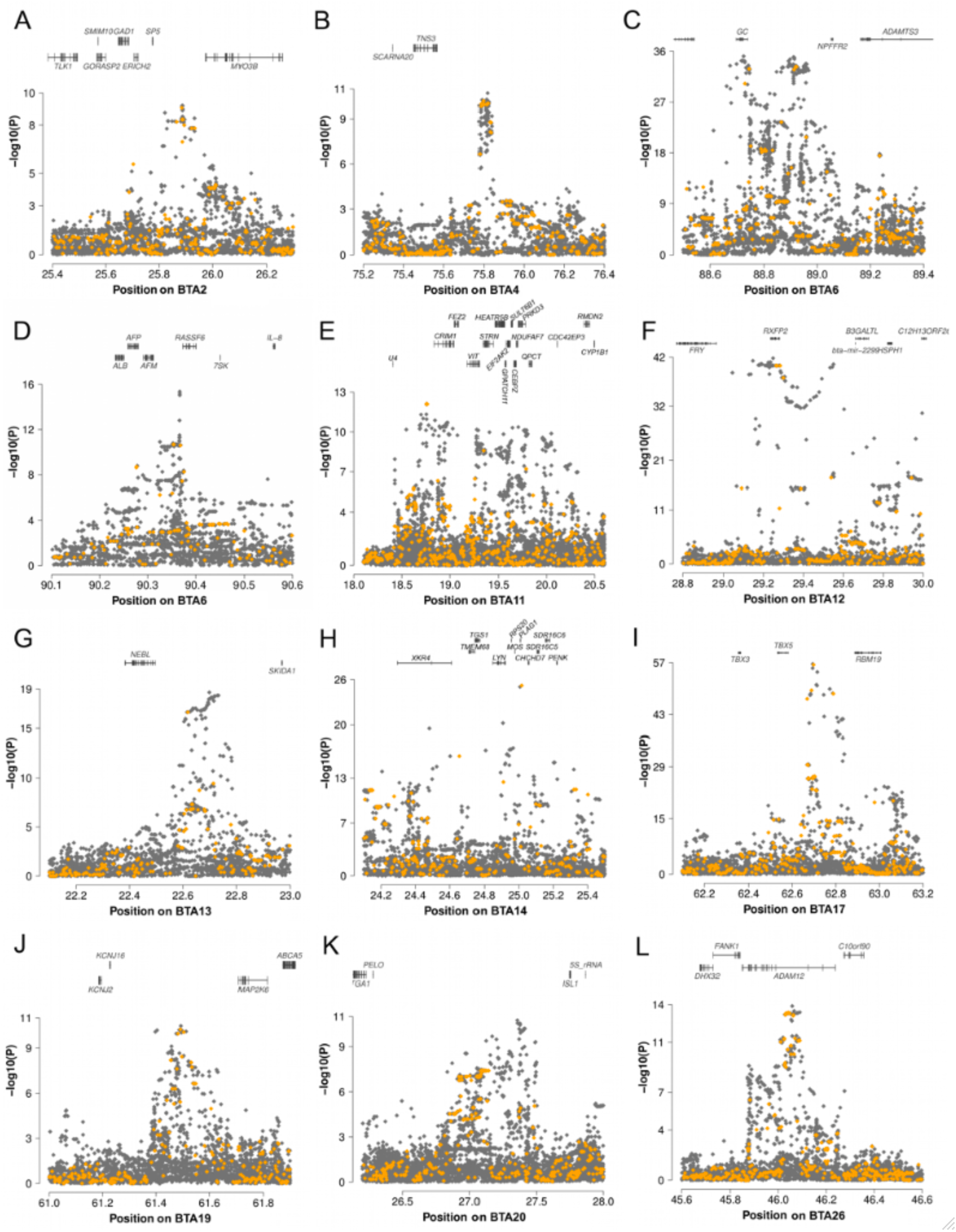
Gene content within twelve QTL regions. Grey and orange dots represent imputed sequence variants and 700K array-derived variants, respectively.

**Additional file 4.**
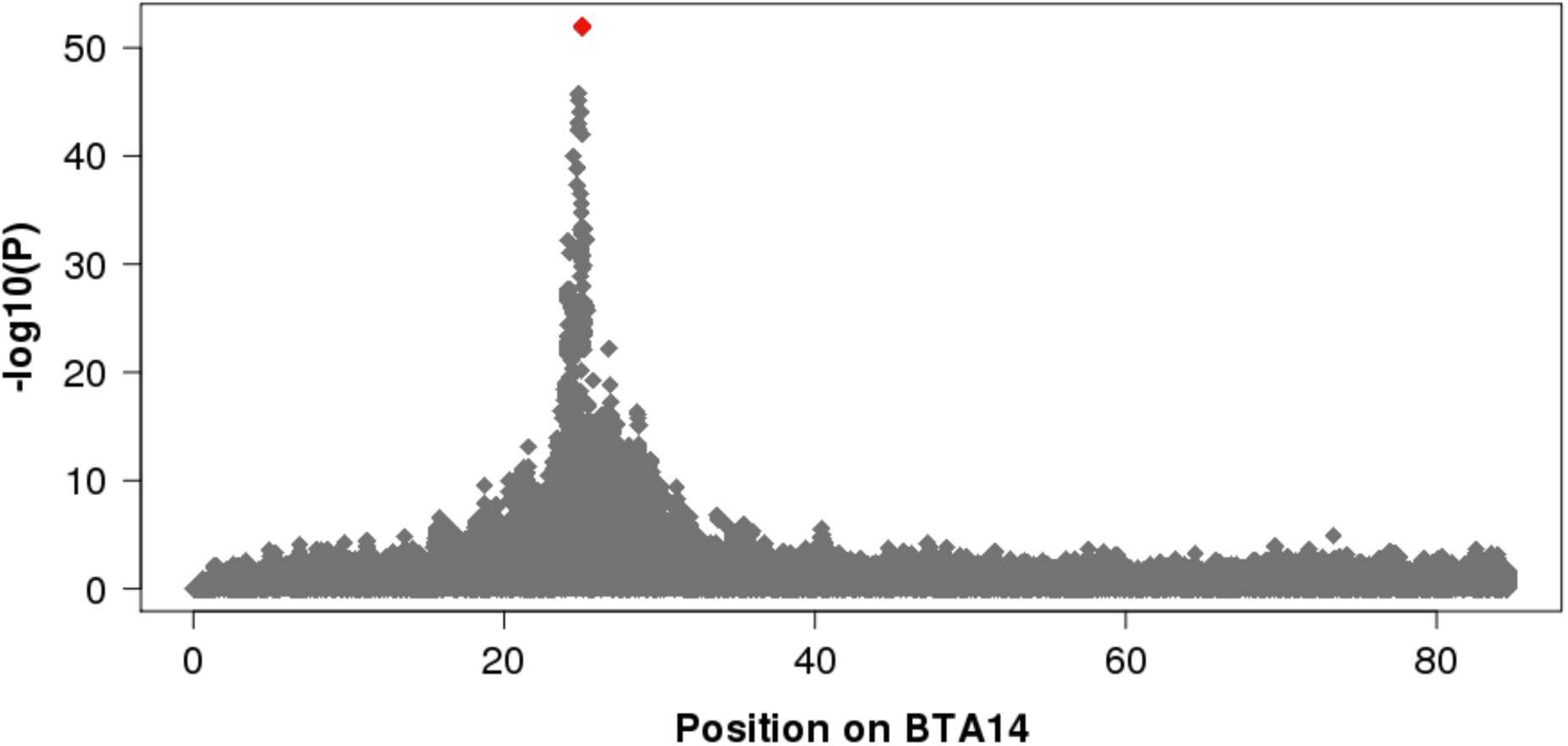
A stature QTL on BTA14. Association of 682 047 imputed sequence variants on BTA14 with height at the sacral bone in 6838 animals. The red diamond represents the most significantly associated variant (BovineHD1400007259).

**Additional file 5.**
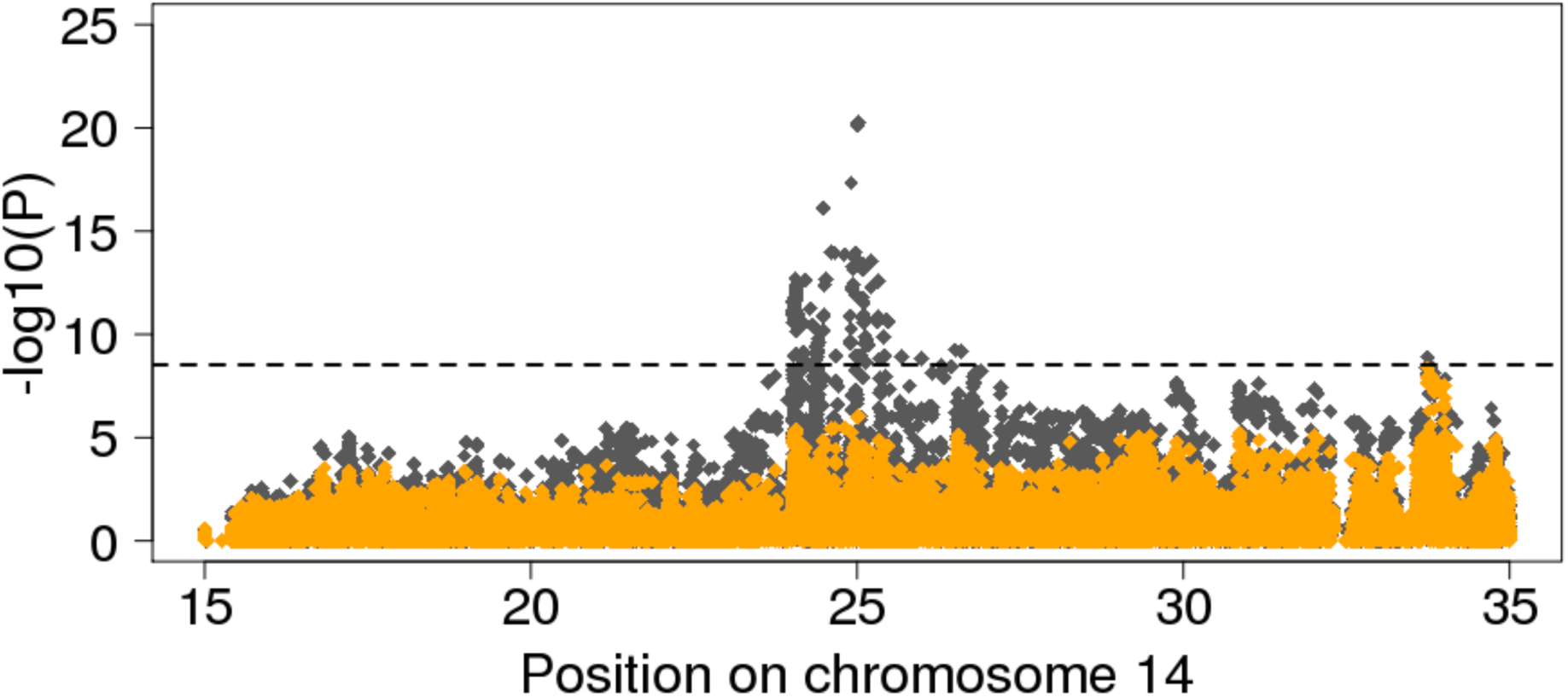
Detailed view of a mammary gland morphology QTL on BTA14. Association of 121 912 imputed sequence variants on BTA14 with udder depth. Grey and orange diamonds represent the −log10(P_SINGLE_)-values from the raw analysis and from the analysis conditional on the DYDs for height at the sacral bone, respectively. The dotted line represents the Bonferroni-corrected significance threshold.

